# Metabolic resistance of the tiger mosquito to pyrethroid insecticides in La Réunion Island likely results from local adaptation

**DOI:** 10.1101/2025.08.06.668844

**Authors:** Tiphaine Bacot, Jean-Marc Bonneville, Vincent Lacroix, Pascal Oberbach, Thierry Gaude, Louis Nadalin, Frederic Laporte, Nausicaa Habchi-Hanriot, Guillaume Dupuy, Cyrille Czeher, Michael C Fontaine, Frederic Boyer, Jean-Philippe David

**Author notes:** Equal contribution (Tiphaine Bacot, Jean-Marc Bonneville).

## Abstract

The resistance of mosquitoes to insecticides is a valuable model system for studying the genetic bases of xenobiotic adaptation in insects. The spread of the Asian tiger mosquito *Aedes albopictus* combined to the massive use of pyrethroid insecticides to limit arbovirus transmission resulted in the rise of resistance in various continents. Here, we investigated the genetic mechanisms underlying the recent adaptation of this mosquito to deltamethrin in La Réunion island. Bioassays confirmed the presence of resistance alleles in field populations. The resistance phenotype was further enhanced in the laboratory following a few generations of controlled selection. Combining whole genome Pool-seq and RNA-seq revealed no evidence of target-site resistance mutations but the over-expression and variant selection of detoxification enzymes associated with pyrethroid metabolism including cytochrome P450s, transferases and ABC-transporters. Among over-expressed detoxification genes, only one was linked to a gene duplication while polymorphism data suggest most of them being trans-regulated. Genome-wide selection signatures revealed a 9 Mb inverted superlocus responding to insecticide selection whose phenotypical importance remains uncertain. Altogether, this study indicates that the multigenic metabolic resistance phenotype observed in this insular territory mainly results from local adaptation. From an applied perspective, this study provides a set of markers to track pyrethroid resistance in the tiger mosquito in the South-West Indian Ocean. As this region is subjected to recurrent arbovirus outbreaks, the additive resistance phenotype that may arise from the introduction of *Kdr* mutations from other territories also calls for improving resistance surveillance at the regional scale.

**Author summary:** While novel vector control strategies are being developed, chemical insecticides remain widely used to control mosquitoes transmitting human diseases such as the Asian tiger mosquito. However, the recurrent use of insecticides resulted in the emergence of resistance which can ultimately affect vector control efficacy. Here, we investigate the genetic bases underlying the resistance of the Asian tiger mosquito to the pyrethroid insecticide deltamethrin in La Réunion island. By combining two complementary genomic approaches, we showed that resistance is mainly caused by an increased insecticide detoxification while classical ‘Knock down resistance’ mutations affecting the target of the insecticide were not detected. We also showed that resistance is underlain by multiple genetic changes spread across the genome, supporting the local selection of resistance rather than the introduction of resistance alleles. Furthermore, we identified a large inverted supergene responding to insecticide selection. This study provides valuable insights into the genetic bases of insecticide resistance, enabling the implementation of molecular makers to improve the tracking of insecticide resistance in this major mosquito vector across the Indian Ocean.

## Introduction

Natural populations experience a variety of human-driven selective pressures, leading to their adaptation through the expression of complex phenotypes (1). Understanding the genetic mechanisms allowing them to adapt to such rapid environmental changes provides key knowledge for populations management (2–4). The evolution of insecticide resistance is a good example of how natural selection can mediate the rapid adaptation of natural populations to anthropogenic pressures as it occurs in a large number of insects (5,6). Beyond understanding the adaptative mechanisms, identifying resistance alleles also contributes to improving management strategies (7–9). However, the frequent co-occurrence of multiple resistance mechanisms and the diversity of genetic events underlying resistant phenotypes often makes the characterization of resistance alleles challenging (5,10).

Among insect vectors, Aedes mosquitoes pose a significant threat to public health because they transmit major arboviral diseases (11,12). Among them, the invasive Asian tiger mosquito Aedes albopictus is of particular importance because its spread contributed to the worldwide re-emergence of arboviruses such as dengue and chikungunya (13). Although significant research efforts are dedicated to develop vaccines targeting these arboviruses (14–16), outbreak prevention still massively relies on insecticide-based vector control (17). Pyrethroid insecticides such as deltamethrin are the most-used insecticides against Aedes mosquitoes (18). However, the adaptation of mosquitoes to such insecticides can ultimately reduce the efficacy of vector control preventive actions (19–22).

Pyrethroids disrupt nerve function by altering the opening kinetics of the voltage-gated sodium channel (VGSC gene: 23,24). Pyrethroid resistance can be the consequence of point mutations affecting the VGSC gene, known as Knock-down resistance (*Kdr*) mutations (target-site resistance); a decreased insecticide penetration mediated by cuticle modification (cuticle resistance); and biodegradation of the insecticide through complex detoxification pathways (metabolic resistance: 10). *Kdr* mutations typically affect a few conserved amino-acid positions in the VGSC protein and are fairly well characterized in *Aedes* mosquitoes (25–28). Conversely, the genetic bases of insecticide detoxification are far less understood. Indeed, the diversity and redundancy of genes underlying xenobiotic detoxification pathways often lead to various multigenic adaptive trajectories depending on the local context (5,10,17,23). In insects, pyrethroid metabolism typically includes an initial phase involving hydroxylation by cytochrome P450s and sometimes ester-bond hydrolysis by carboxylesterases, followed by the conjugation of hydrophilic groups by glutathione S-transferases (GSTs) and UDP-glycosyltransferases (UDPGTs), and the excretion of the insecticide itself or its metabolites by ABC-transporters (29–31). Molecular changes underlying metabolic resistance to pyrethroids frequently include the over-expression of detoxification enzymes and transporters mediated by cis-or trans-regulation or by gene duplication although the selection of enzyme/transporter variants showing an enhanced activity toward the insecticide has also been described (6,32–35).

The recent spread of *Ae. albopictus* to all continents and its ecological shift to more urbanized areas (12) led to its increased exposure to pyrethroid insecticides and ultimately to the selection of resistance. Pyrethroid resistance was first reported in South-East Asia more than a decade ago (36–39) followed by other Asian regions (40–42). In these regions the resistance phenotype was often related to the presence of various *Kdr* mutation haplotypes (40,43,44), although metabolic resistance was also detected (45–47). Resistance was then detected in Europe following the introduction of *Kdr* mutations from Asia (48–50). Resistance was also detected in Africa in association with *Kdr* mutations and increased detoxification though the underlying resistance alleles were not characterized (51–53).

In the Indian Ocean, *Ae. albopictus* was likely imported from Asia in La Réunion island during the 17^th^ and the 18^th^ centuries though other introductions likely occurred later (54). This mosquito was associated to the chikungunya outbreak that occurred in 2005-2006 (55,56) and is the main vector of recurrent dengue/chikungunya outbreaks since then (57). In La Réunion island, the recurrent use of deltamethrin for vector control since 2006 ultimately resulted in the emergence of resistance as detected by public health authorities in 2017 (58). However, the mechanisms underlying the resistance phenotype have not been thoroughly investigated since then. A single heterozygote for the V1016G *Kdr* mutation was observed from mosquitoes collected in La Réunion in 2016 as part of a worldwide *Kdr* screening study (59). However, the circulation of *Kdr* mutation in the island or neighbouring territories was not confirmed after this observation.

In this context, the present work aims at deciphering the genetic bases of the deltamethrin resistance phenotype observed in *Ae. albopictus* populations in La Réunion island. This was achieved by adopting an experimental design combining field populations and field-derived lines enriched or not for resistance alleles by controlled selection. At the molecular level, an integrated genomic approach combining RNA-seq and whole genome Pool-seq was applied to pinpoint resistance alleles and their underlying genomic events. The evolutionary scenario leading to the emergence of resistance in La Réunion island is then discussed in regards of the worldwide distribution of resistance alleles and the local demographic context. While providing a set of novel molecular markers to track insecticide resistance in this mosquito vector, the present study contributes to anticipate the emergence of insecticide resistance in the South-West Indian Ocean region and its consequences for vector control strategies.

## Results

### Deltamethrin resistance alleles circulate in La Réunion island

In order to improve the identification of deltamethrin resistance alleles, an experimental design combining field populations and the controlled selection of a composite line was used (**Fig. 1A**, see methods). Comparative deltamethrin bioassays performed on three field populations confirmed the presence of resistance alleles in La Réunion (**Fig. 1B**). The three field populations (Saint-Paul, Saint-Louis and Sainte-Suzanne) showed a significant decreased mortality to deltamethrin compared with the susceptible SRun reference line. As expected, the composite line maintained without deltamethrin selection for 6 generations (NS line) showed a resistance level similar to field populations. Selecting the composite line for 5 generations with deltamethrin (Sel line) resulted in its increased resistance as compared to the NS line, suggesting that resistance alleles are not fixed in the field and have been further selected by controlled-selection in the laboratory.

**Fig 1.**
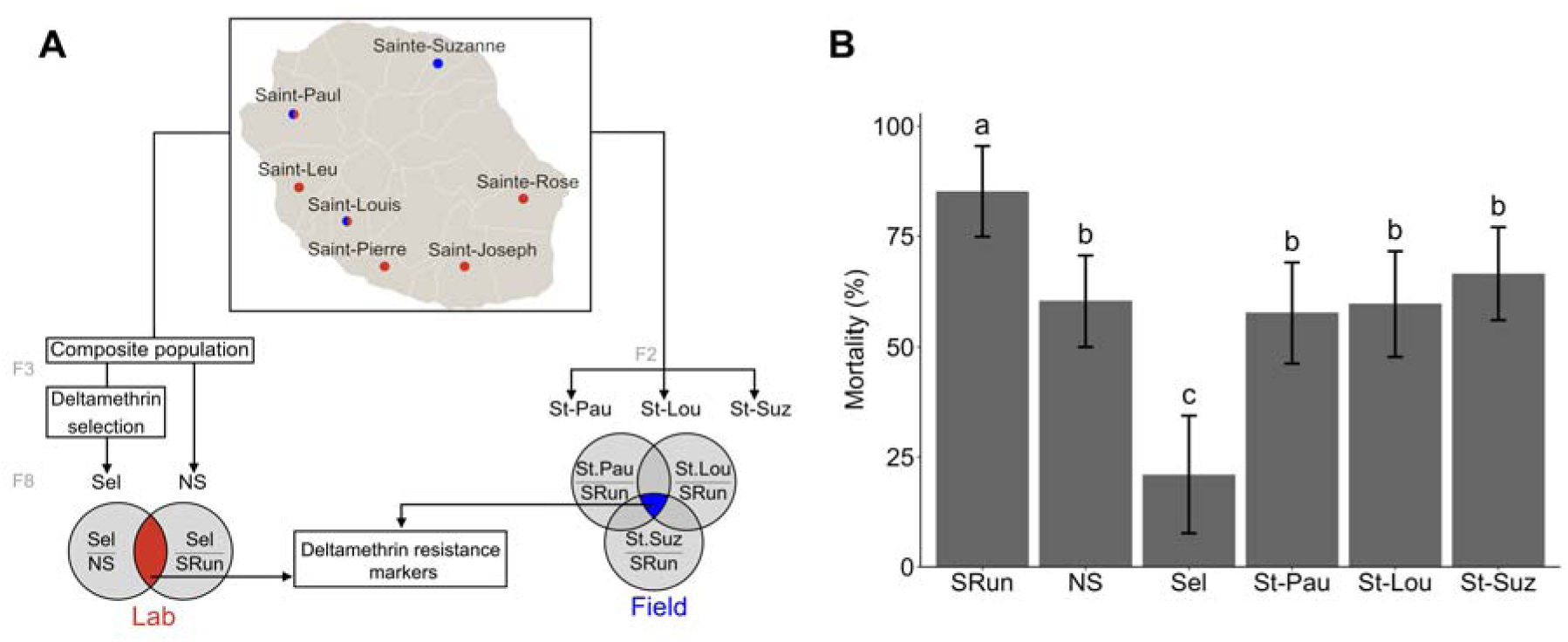
Study design and deltamethrin resistance levels. (A) Sampling sites and study design. The map shows the location of populations used to create the composite population from La Réunion island (red dots) and field populations (blue dots) used for analyses. For each line/population, the generation used is indicated. Venn diagrams illustrate the pairwise comparisons made for the identification of deltamethrin resistance markers (see methods). (B) Deltamethrin resistance levels. Mean mortality rates were obtained from standard WHO test tube assays using 0.03% deltamethrin impregnated papers. Error bars denote SD. Distinct letters indicate significantly distinct mortality rates (Conover-Iman test; N=9; *p value* ≤ 0.05). Sel: deltamethrin selected line; NS: non-selected line; St-Pau: Saint-Paul; St-Lou: Saint-Louis; St-Suz: Sainte-Suzanne; SRun: susceptible reference line from La Réunion.

### The resistance phenotype is not associated with target-site mutations

Whole genome pool-seq data from all lines were used to screen for *Kdr* mutations affecting the VGSC gene encoding the biochemical target of deltamethrin. Among the 132 polymorphisms detected in coding regions, six were non-synonymous (**S1 File**), but none matched the position of *Kdr* mutations previously associated with pyrethroid resistance in *Ae. albopictus* (i.e. V410L, L982Y, S989P, I1011M, V1016G/I, I1532T, and F1534C:, 28). In addition, these six non-synonymous variants are unlikely to contribute to deltamethrin resistance as they do not show a frequency pattern consistent with deltamethrin resistance across all lines. Furthermore, they were located in cytosolic regions being distant from the protein domains carrying known *kdr* mutations. Altogether, this indicate that the detected protein variants are most likely neutral, and that *Kdr* mutation do not contribute to deltamethrin resistance in La Reunion island.

### Resistance involves the over-transcription of detoxification enzymes

RNA-seq performed on adult females enabled the detection of 13,918 genes passing coverage threshold which included 697 resistance candidate genes. Differential transcription analysis identified 48 genes differentially transcribed in association with resistance when considering both lab and field comparisons (**Fig. 2A and S2 Table**). The 12 under-transcribed genes include a single candidate encoding an ABC-transporter (LOC109429564). Conversely, the 36 over-transcribed genes included 11 candidates, representing a significant enrichment compared with the global gene set (from 5% to 30.5%, Χ test *p* < 0.05). All of them belong to three gene families classically involved in pyrethroid metabolism (P450s, transferases and ABC-transporters), leading to a significant enrichment of detoxification genes (from 54% to 100%, Χ test p < 0.05; **Fig. 2B**). All over-transcribed candidate genes were located on chromosomes 1 and 2, some of them being clustered together (**Fig. 2C**). Several of them show high protein homology with genes previously involved in pyrethroid resistance in *Aedes aegypti or Anopheles gambiae*. The P450s *CYP6d5* (LOC115259486) was previously found over-transcribed in resistant *Ae. albopictus* populations from China (same gene named CYP6A8 in 47) and shows high homology with members of the CYP6Z subfamily previously involved in pyrethroid metabolism (60–62). The P450 CYP6a14 (LOC109421559) shows high homology with members of the CYP6M subfamily validated as pyrethroid metabolizers in Anopheles mosquitoes (62,63). Similarly, GSTl-1 (LOC109412993) and GST4-like (LOC109413318) showed high homology with *theta*– and *delta*-GSTs previously associated with pyrethroid in mosquitoes (45,64–67). Finally, the two ABC-transporters (LOC109407008 and LOC109429691) showed a high homology with the ABCC subfamily previously associated with pyrethroid resistance (68–73). Altogether, these results indicate that in absence of *Kdr* mutations, the deltamethrin resistance phenotype observed in La Réunion island is associated with an over-expression of detoxification enzymes likely involved in insecticide metabolism.

**Fig 2.**
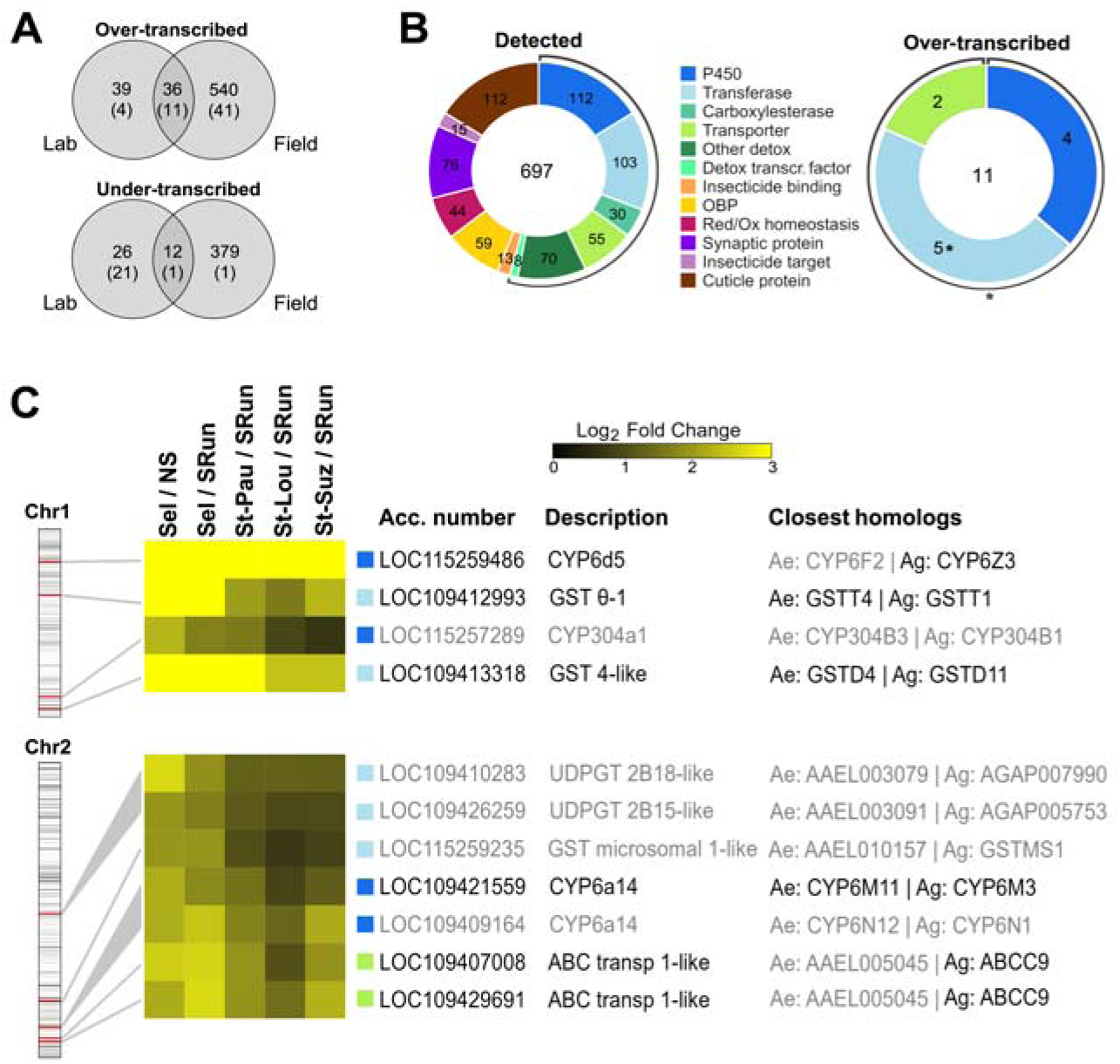
Genes identified as differentially transcribed in association with deltamethrin resistance. Only genes showing consistent transcription level variations in both Lab and Field comparisons were considered associated with resistance (see methods). (A) Venn diagrams showing the number of genes being differentially transcribed in association with resistance. Numbers within brackets stand for candidate genes. (B) The left donut-chart describes all candidates genes detected by RNA-seq and their respective gene families. The right donut-chart shows candidate genes associated with resistance. Black circular arcs indicate gene families classically associated with detoxification. Stars indicate a significant enrichment from ‘detected’ to ‘resistance associated’ candidate genes (Χ^2^ test, *p* value ≤ 0.05). (C) Candidate genes over-transcribed in association with deltamethrin resistance. Log_2_ fold changes obtained from each comparison are shown on the heatmap (scale from 0 to 3, see S2 table for exact values). Colour squares refers to candidate gene families. Chromosomal positions are shown on the left with the density of detected candidate genes shown as grey intensity. For each gene, NCBI accession number and description are provided together with closest protein homologs in *Ae. aegypti* and *An. gambiae*. Genes in bold are those from which close orthologs were previously associated with pyrethroid resistance. CYP: cytochrome P450; GST: glutathione S-transferase; UDPGT: UDP-glycosyltransferase; ABC transp: ABC transporter; Sel: selected line; NS: non-selected line; St-Pau: Saint-Paul; St-Lou: Saint-Louis; St-Suz: Sainte-Suzanne; SRun: susceptible reference line from La Réunion.

### Resistance is likely not associated with alternative splicing

Using a *de novo* RNA-seq assembly strategy, we searched for alternative splicing events associated with resistance. This analysis identified 166 events associated with resistance affecting 163 genes (**S3 Table**). This included 8 alternative splicing events, 4 large intron retention events (> 50 bp) though none of them affected candidate genes. Such analysis also identified 161 short genomic indel events (≤ 50 bp) among with two affected candidate genes: the transcription factor HR96 LOC109416347 (6 bp indel) and the peroxiredoxin LOC109415663 (3 bp indel). Altogether, these data indicate that resistance in la Réunion does not strongly rely on alternative splicing events.

### An over-transcribed P450 is likely affected by a genomic duplication

As over-transcription can be caused by an increased copy number, whole genome pool-seq data were used to identify genes affected by copy number variations (CNV) in association with resistance based on their increased exonic read coverages (see Methods). This analysis identified 72 genes showing an increased coverage associated with resistance. Five of them were also over-transcribed in resistant lines including the P450 *CYP6d5* (**Fig. 3A and 3B; S4 Table**). Comparing normalized coverage profiles at the *CYP6d5 locus* revealed an increased coverage in resistant lines, which supports the occurrence of a ∼60 Kb duplication (**Fig. 3C**). The putative breakpoints were poorly covered, likely due to the presence of transposable elements which were annotated as a LTR element on the left and a Line/R1 element on the right.

**Fig 3.**
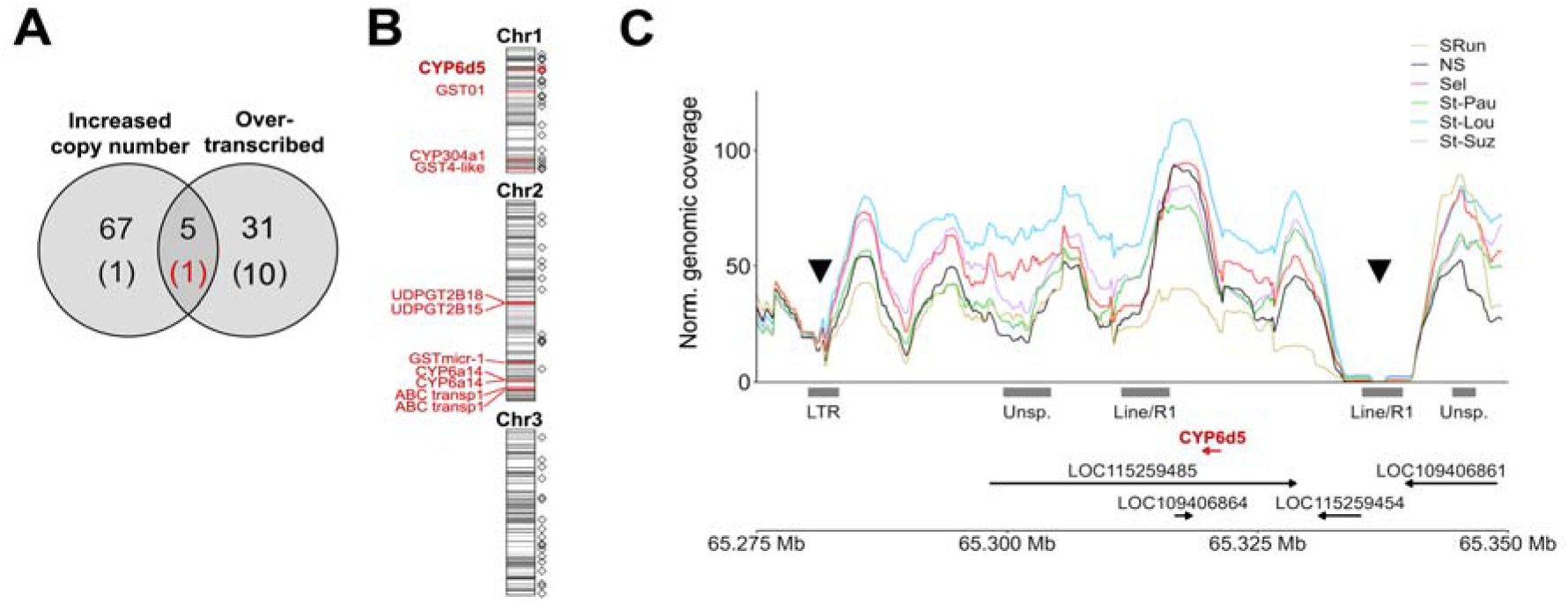
Gene copy number variant analysis. (A) Genes showing an increased copy number (CNV) in association with deltamethrin resistance in regards of those identified as overtranscribed. Numbers within brackets refer to candidate genes. (B) Distribution of gene copy number variants along chromosomes. Diamonds show the chromosomal position of the 72 genes showing an increased CNV associated with resistance. The red diamond refers to the candidate gene *CYP6d5* (LOC115259486). Over-transcribed candidate genes are named and shown as red marks in respect to candidate genes density (grey intensity). (C) Normalized read coverage profiles observed at the vicinity of *CYP6d5*. Normalized coverage profiles were computed for each line from whole genome pool-seq reads and smoothed using a 5 kb sliding window. The probable range of the duplicated region is delimited by black triangles. The *CYP6d5* gene is shown in red. Repeat masker annotation is shown as grey boxes. Genomic coordinates refer to AALBF3.v2 assembly. Sel: selected line; NS: non-selected line; St-Pau: Saint-Paul; St-Lou: Saint-Louis; St-Suz: Sainte-Suzanne; SRun: susceptible reference line from La Réunion.

### Resistance is related to the selection of detoxification enzyme variants

Whole genome Pool-seq data were used to look for non-synonymous variants associated with resistance based on their frequency variation across resistant and susceptible lines. This analysis detected 197,403 non-synonymous variants passing coverage threshold with 11,679 of them affecting candidate genes (**Fig. 4A**). Among the 455 non-synonymous variants associated with resistance, 36 of them affected 27 distinct candidate genes **(Fig. 4B, S5 Table)**. A significant enrichment of candidate genes involved in detoxification was observed, mainly driven by a strong enrichment of P450s (from 15% to 33.3%, Χ^2^ test *p* < 0.02). The 36 variants affecting candidate genes were spread over the three chromosomes (**Fig. 4C**). None of them affected over-transcribed genes though some were located in the same gene cluster. This was the case for the P450 *CYP6a8* variant located within a large CYP6 cluster which also include the two over-transcribed *CYP6a14* genes. Four major loci gathering multiple variants were identified in chromosome 3. The first one carried 4 variants affecting a *CYP4v2* and an ABC transporter. The second one included 9 variants affecting 6 P450s belonging to a cluster of 18 *CYP4C* genes. The last two gathered various candidate genes including a *CYP9f2* gene and an epoxide. Altogether, this analysis indicates that the deltamethrin resistance phenotype observed in La Réunion is not only caused by the overtranscription of detoxification genes but also by the selection of protein variants likely related to insecticide metabolism.

**Fig 4.**
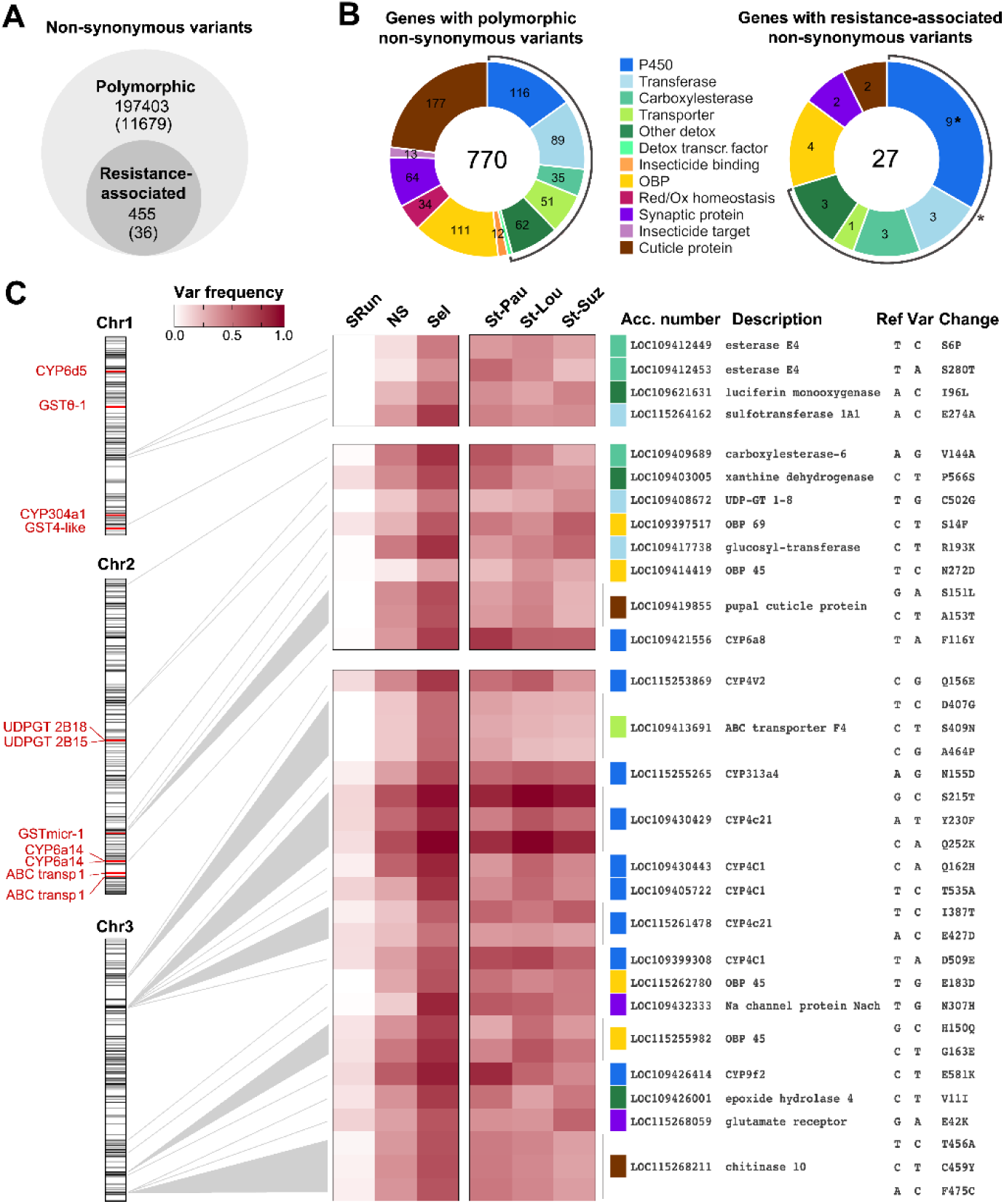
Protein variants associated with resistance. (A) Overview of non-synonymous variants identified. Numbers within brackets refer to variants affecting candidate genes. Variants associated with resistance were identified based on allele frequency variations across resistant and susceptible lines (see Methods) (B) The left donut-chart describes all candidate genes from which polymorphic non-synonymous variants were detected. The right donut-chart described those associated with resistance. Black circular arcs highlight gene families associated with detoxification. Stars indicate a significant enrichment (Χ^2^ test, *p* value ≤ 0.05). (C) Candidate genes with non-synonymous variants associated with resistance. The heat map shows the frequency of the variant allele across all lines. The chromosomal position of each variant is shown on the left in respect to over-transcribed candidate genes (red marks) and candidate gene density (grey intensity). For each variant, gene family (colour square), gene accession number, gene description, nucleotide and amino-acid change are indicated. CYP: cytochrome P450; GST: glutathione S-transferase; UDPGT: UDP-glycosyltransferase; ABC transp: ABC transporter; Sel: selected line; NS: non-selected line; St-Pau: Saint-Paul; St-Lou: Saint-Louis; St-Suz: Sainte-Suzanne; SRun: susceptible reference line from La Réunion.

### Over-transcribed detoxification genes are not under direct selection

Exonic polymorphism data were then used to look for gene-based selection signatures across the genome between the Sel and NS lines based from allele frequency differentiation indexes (see Methods). Among the 11 over-transcribed candidate genes, only the P450 *CYP304c1* and the microsomal *GST-1* showed high differentiation indexes, which supports their cis-regulation. Other over-transcribed candidate genes were below the baseline, consistent with their trans-regulation (**Fig. 5A**). Conversely, most candidate genes carrying protein variants associated with resistance showed high gene differentiation indexes, supporting selection acting on the same *loci*. Among candidate genes, those showing the highest differentiation indexes included two carboxylesterases *E4-like* on chromosome 1, two UDPGTs and the *CYP6a8* P450 on chromosome 2, and the ABC transporter *F4* and the two *CYP4c1* P450s on chromosome 3. Overall, though genetic drift may have occurred between the Sel and NS lines, the high differentiation indexes observed across the three chromosomes support the hypothesis of a multigenic adaptation based on standing genetic variation in La Réunion island (**Fig. 5B**).

**Fig 5.**
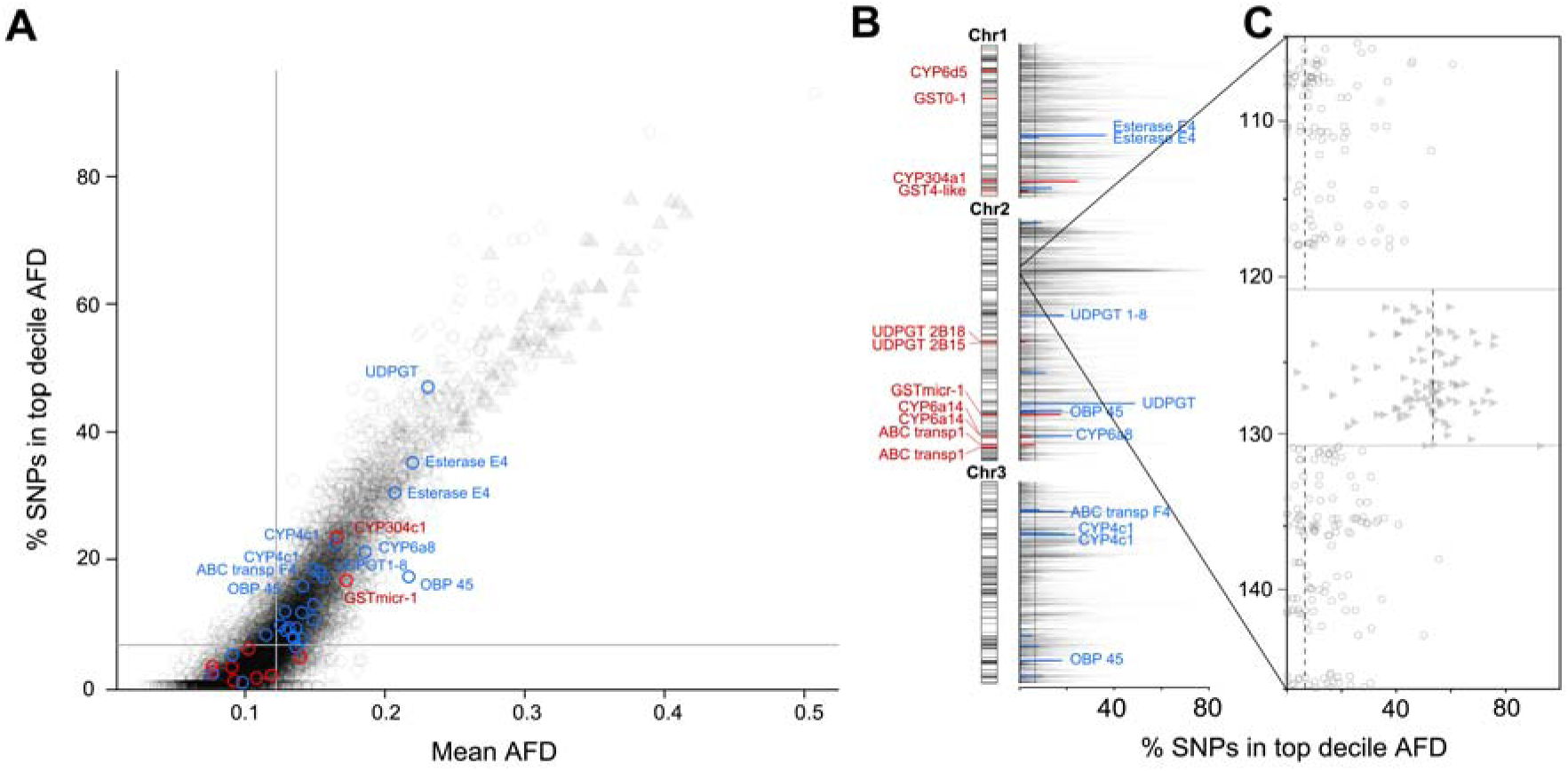
Genome-wide allele frequency differentiation. (A) The plot shows two genetic differentiation indexes for the Sel/NS line comparison: the mean Allele Frequency Difference per gene (mean AFD, absolute value) and the proportion (%) of SNPs per gene showing an AFD within the top decile of the global distribution (% SNPs in top decile AFD). These differentiation indexes were computed for the 10,192 protein-coding genes carrying > 20 bi-allelic substitutions (see Methods). Median values are shown as grey lines. Candidate genes within top quartiles for both differentiation indexes are annotated with those differentially transcribed in association with resistance in red and those carrying protein variants associated with resistance in blue. Triangles denotes genes carried by the genomic inversion identified in chromosome 2. (B) Genome wide overview of the ‘% SNPs in top decile AFD’ index for the Sel/NS comparison. The dashed line stands for the median of the differentiation index across all genes. The chromosomal position of candidate genes associated with resistance is shown together with candidate gene density (grey intensity). (C) Focus on the genomic region carrying the 9 Mb inversion associated with resistance. The ‘% SNPs in top decil AFD’ index is shown for all genes. Horizontal grey lines delimit the inverted region with breakpoints at Chr2:120,519,062-130178023 in AALBF3.v2 assembly (Chr2:405,097,032-416,198,432 in AALBF5 assembly). Vertical dashed lines show the median of the ‘% SNPs in top decile AFD’ index within and outside the inverted region. The genomic scale is in Mb respective to AALBF3v2.

### An inverted superlocus is under deltamethrin selection in La Réunion

This analysis also identified a hundred genes located within an intriguing region of chromosome 2 displaying very high differentiation levels (**Fig. 5C and S6 Table**). This large 9.6 Mb *locus* showed a strong selection signal in the Sel/NS comparison but also in other ‘resistant vs susceptible’ comparisons coherent with a low recombination rate. Indeed, a quarter of the 3141 non-synonymous polymorphisms detected in this region are above the whole genome top centile for Sel/NS differentiation indexes. Examining reads topology identified ‘one mate swap’ read pairs and split reads supporting the presence of a large genomic inversion whose breakpoints were 120,519,062 and 130,178,023. The partial mapping of these reads at the breakpoints together with the unmapped fragments matching to multiple genomic locations supported the flanking of this inversion by two transposable elements. The presence of this inversion at low frequency in the susceptible reference strain SRun also suggests its pre-existence prior to its selection by deltamethrin in La Réunion. Among the 107 genes located within the inverted region, only four belong to candidate genes including two ABC-transporters, a cytochrome b5-like and a cuticle protein. However, none of them were differentially transcribed, affected by CNV, or carried non-synonymous variants associated with deltamethrin resistance. Among other genes located within the inversion, none showed a clear association with a known resistance mechanism. Fifteen genes encoded known or putative (*i*.e. zing finger proteins) transcription factors although none were previously associated with detoxification enzymes regulation. Several polymorphisms showing the highest Sel/NS differentiation affected the uncharacterized gene LOC115268503. Protein homology suggest that this gene encodes a S/T protein phosphatase with ankyrin motifs. As the voltage-gated sodium channel targeted by pyrethroids can be regulated by phosphorylation and interact with ankyrins (74,75), this gene is an attractive candidate to explain the selection of this large inversion by deltamethrin.

## Discussion

### Metabolic resistance likely emerged from local adaptation in La Réunion island

Bioassays performed on field mosquito populations from La Réunion island revealed a significant deltamethrin resistance phenotype, confirming the rise of resistance observed since 2017 (58). Considering that resistance may have remained undetected for some times, the emergence of resistance may have begun a few years after deltamethrin was adopted for vector control following the 2005-2006 chikungunya outbreak. The resistance level of the non-selected composite line was similar to those observed in the field, suggesting that the sampling design was efficient at capturing resistance alleles circulating in the island. Indeed, the ancient colonization of this insular territory by *Ae. albopictus* was linked to a high local genetic diversity with significant gene flows between populations (76–78).

More than 30 non-synonymous polymorphisms affecting the VGSC targeted by deltamethrin have been identified in *Ae. albopictus* worldwide but only a few were consistently associated with pyrethroid resistance (23,26,28,79: V1016G, I1532T and F1534S/C/L). Despite the sequencing of large mosquito pools at high coverage, none of these *Kdr* mutations was detected in La Réunion island, either in field populations or in composite laboratory lines. Considering the recurrent usage of deltamethrin since 2005, this supports the absence of *Kdr* mutations in La Réunion at the time of the study. As *Kdr* mutations have already spread across multiple continents including Asia, Europe and Africa (26,28), such result also confirm that limited gene flow occurred from these regions. Instead, a multigenic metabolic resistance phenotype was observed, involving detoxification genes belonging to families involved in pyrethroid metabolism such as cytochrome P450s, glutathione S-transferases, UDP-glycosyl transferases, and ABC-transporters (31,80). Elevated detoxification enzyme activities have been linked to deltamethrin resistance in *Ae. albopictus* populations from various regions (46,81–84).

Transcriptomic screenings revealed the over-transcription of the same detoxification gene families in *Ae. albopictus* resistant populations from Asia with some P450s being functionally validated (44,45,47). Among the detoxification genes identified in La Réunion, only the P450 (*CYP6d5*) was previously identified as contributing to deltamethrin resistance in *Ae. albopictus* (same CYP gene named CYP6A8: 47). Yet, several of them showed high protein homology with known pyrethroid resistance genes identified in other mosquito species. This includes two P450s showing high homology with CYP6Ms/CYP6Zs subfamilies, two GSTs likely belonging to GST-theta and –delta families and two ABC-transporters likely belonging to the ABCC family. Although functional validation is needed to clarify their importance in the overall phenotype, the emerging picture is the independent selection of detoxification genes in Asia and La Réunion. Such hypothesis is also supported by the absence of *Kdr* mutations previously detected in Asia in La Réunion island. Such parallel evolution of detoxification genes under pyrethroid selection is probable considering the functional redundancy observed within these gene families whose members are often organized in clusters of genes sharing a high homology (10,85). Altogether, while the introgression of metabolic resistance alleles cannot be ruled out, the genetic isolation of La Réunion island together with the absence of *Kdr* mutations and the specific nature of the genes identified support their local selection from standing genetic variation.

### Multiple genetic events underly the resistance phenotype

Gene duplications mediating the over-expression of detoxification genes have been frequently associated with the rapid adaptation of mosquitoes to insecticides (32,35,86,87). Here, the over-transcribed P450 *CYP6d5* was linked to the presence of a ∼100 Kb genomic duplication. Although our short-read Pool-seq data did not allow to fully resolve the genomic architecture of the duplicated locus, its flanking by transposable elements (TEs) was not surprising as TE-movements are often associated with genomic duplication events (86,88), especially in TE-rich genomes such as Aedes mosquitoes (89–91). No such duplication was identified for the other over-transcribed detoxification genes, implying that their over-transcription is rather mediated by cis-or trans-regulation. In this regard, the nuclear receptors CncC/Maf and AhR-ARNT considered as key regulators of insect detoxification genes (92) were not differentially transcribed nor affected by protein variants in association with resistance. Nevertheless, a two-codons deletion variant affecting the transcription factor HR96 showed an increased frequency in resistant lines. The contribution of this transcription factor to the regulation of detoxification enzymes has been demonstrated in spider mites (93,94). Although this requires further validation, this finding supports the potential role of HR96 in the regulation of detoxification enzymes in *Ae. albopictus*.

As metabolic resistance may also be the consequence of the selection of proteins variants showing a higher activity toward the insecticide (10), polymorphism data were screened for non-synonymous variants associated with resistance. Such analysis identified multiple detoxification protein variants with P450s being over-represented. Yet this gene set showed no overlap with over-transcribed genes, suggesting that these two adaptive mechanisms contribute to the multigenic resistance phenotype observed in La Réunion island. Indeed, in a context of local adaptation, it is expected that selection retains any polymorphism that contribute to resistance, affecting both gene expression and protein function. Of particular interest was the selection of multiple P450 variants belonging to a cluster of 18 *CYP4C*-like genes on chromosome 3. Indeed, while *CYP4*s are not considered as classical xenobiotic metabolizers in mosquitoes as opposed to CYP6s, CYP9s, some of them have been shown to be over-transcribed in association with pyrethroid resistance in *Ae. aegypti* (95), suggesting their possible contribution to deltamethrin metabolism. Similarly, the identification of two carboxylesterase variants associated with resistance may indicate that these enzymes can still be active on deltamethrin or its metabolites, even though the cyano group of type II pyrethroids like deltamethrin is thought to protect the ester bond from a direct hydrolysis (96).

Polymorphism data were then used to look for genes under positive selection in association with resistance. As expected, a good proportion of detoxification protein variants associated with resistance showed a positive selection signal at the gene level. On the other hand, apart from one P450 and one GST, most over-transcribed detoxification genes showed poor selection signals, supporting their trans-regulation. Unexpectedly, such analysis also identified a large ∼9 Mb genomic inversion on chromosome 2 behaving as a supergene under deltamethrin selection. Genomic inversions can play a key role in local adaptation as demonstrated in various biological models (97–100). Whether this genomic event is specific to the South-West Indian Ocean or more widely distributed remains to be investigated. In that concern, the identification of the inversion breakpoints will facilitate its high-throughput genotyping by PCR. The hundred genes carried by the inverted locus included four candidates but none of them were directly associated with deltamethrin resistance, suggesting that they do not drive its selection. Considering the strong genetic linkage of the inverted haplotype, deciphering which gene(s) drive the selection of this supergene is not at reach from the current data set and requires functional approaches.

### A warning bell for the South-West Indian Ocean

The French territory of La Réunion island is a hotspot of arbovirus transmission in the South West Indian Ocean in line with a high human density and the presence of *Ae. albopictus* (57). Such epidemiological context led to the recurrent use of deltamethrin for vector control since the first Chikungunya outbreak of 2005-2006. In a context of genetic isolation, this repeated insecticide pressure resulted in the local selection of a metabolic resistance phenotype underlain by various genetic events. Such evolutionary scenario is supported by the multigenic nature of the phenotype together with the striking absence of *Kdr* mutations, which are already widely distributed in this species (26,28). At the time of the study, the resistance phenotype observed in the field was moderate and can be further increased following a few generations of deltamethrin exposure, suggesting that the selection of resistance alleles is undergoing in La Réunion and might be counterbalanced with fitness costs. As the selection of this multigenic resistance phenotype may affect various mosquito life traits including vector competence (7,101), precising its impact on population dynamics and viral transmission is of interest.

Considering that the selection process was undergoing by the time of the study, the intense vector control activity associated with recent chikungunya and dengue outbreaks may have resulted in an increased frequency of resistance alleles in La Réunion and their dispersal to other neighbouring territories of the South-West Indian Ocean. In addition, although *Kdr* mutations were not detected by the time of the study, their introduction from Europe, Asia or Africa should be considered as a major risk for the region. Indeed, their introduction in populations already carrying metabolic alleles is expected to lead to an additive resistance phenotype that may impact vector control efficacy, as shown in *Ae. aegypti* (19,21). In this context, the present study calls for an improved surveillance of pyrethroid resistance in this region. Ideally, this surveillance should combine annual bioassays on sentinel populations and the tracking of key metabolic resistance alleles identified here together with known *Kdr* mutations.

## Methods

### Mosquito lines and experimental design

All mosquito lines were maintained under standard insectary conditions (27 ± 2°C, 70 ± 10 % relative humidity and light/dark cycle 14h:10h). Populations size were maintained over 1000 individuals to reduce the effect of genetic drift. Larvae were reared in tap water and fed with hay pellets. Adults were maintained in mesh cages and fed on filter papers impregnated with a 10 % honey solution. Blood feeding was conducted on mice to generate eggs of the next generation. Three days-old non-blood-fed adult females were used for bioassays and molecular work.

The experimental design combined different mosquito populations all originating from La Réunion island (**Fig. 1A**). First, a composite population from la Réunion island was created by pooling >500 larvae from six field populations collected in 2018-2019 using a minimum of three ovitraps per population. After two generations of genetic mixing (F1 to F3), the composite population was split in two separated lines (N>1000 in each line): the first being maintained without insecticide selection (NS line) and the other being selected with deltamethrin for five successive generations (Sel line). Selection was conducted on males and females separately prior mating as follows. Batches of 25 non-blood-fed 3-5 days-old adults were exposed to filter papers impregnated with 0.03% deltamethrin using WHO test tubes. Exposure times were adjusted from 30 min to 1h to reach 40-60% mortality for each sex. Surviving males and females were then pooled in a single mesh cage and allowed to reproduce. Females were blood fed and allowed to lay eggs to generate the next generation. Three field populations sampled in 2021 from the localities of St-Paul, St-Louis and St-Suzanne were also used. Finally, the laboratory susceptible strain SRun colonized from La Réunion island in the 2000s was used as reference line. For molecular analyses, genes and polymorphisms associated with resistance were identified based on the five pairwise comparisons: ‘Sel vs NS’; ‘Sel vs SRun’ and ‘each field population vs SRun’. Only genes and polymorphisms showing consistent variations (in the same direction) for all comparisons were retained.

### Bioassays

Comparative deltamethrin bioassays were performed on 3-5 days-old adult females following WHO mosquito testing guidelines (102) using test tubes equipped with home-made insecticide papers impregnated with 0.03% deltamethrin. Each bioassay was conducted using 9 replicates of 25 females and an exposure time of 1h. Such insecticide dose resulted in a mortality >80% in the susceptible reference line SRun. After insecticide exposure, mosquitoes were allowed to recover for 24h with a cotton bowl impregnated with a 10% honey solution before mortality recording.

### Reference genome and candidate genes

Genomic analyses were performed using the version AALBF3 of *Ae. albopictus* reference genome, which was the latest available assembly at the time of this study (103). As this assembly was made from 574 ordered scaffolds, the successive scaffolds were assembled into pseudo-chromosomes with 50N spacers (hereafter referred to as AALBF3.v2 assembly). A gff3 annotation file based on AALBF3.v2 assembly was also created based on AALBF3 annotation. The genomic coordinates of each scaffold on AALBF3.v2 assembly is provided in **S7 Table**. Among the 22,435 features annotated in AALBF3.v2, only the 17,342 genes and pseudogenes carried by the three nuclear chromosomes were retained for genomic analyses. A subset of 953 genes belonging to 12 different families possibly involved in insecticide resistance was defined. These candidate genes included: insecticide targets; synaptic proteins; detoxification enzymes sensu lato; known detoxification transcription factors; ABC-transporters; insecticide binding proteins; cuticle proteins; red/ox enzymes; odorant binding proteins (**S8 Table**).

### RNA-sequencing

For each line, four RNA-seq libraries were prepared from distinct batches of 25 five days-old non-blood-fed females. Total RNA was extracted using TRIzol® (ThermoFisher Scientific) according to manufacturer’s instructions. RNA extracts were quantified with a Nanodrop One (ThermoFisher Scientific) and diluted to meet sequencing requirements. Sequencing was performed by the BGI Hongkong Tech Solution NGS Lab. Sample quality control was conducted on an Agilent 2100 and libraries were prepared from 10 ng total RNA following the DNB-seq Eukaryotic Strand-specific mRNA library protocol. Briefly, mRNAs were enriched with poly-A tails using oligodT beads and fragmented to an average size of 200-400 bp before being reverse transcribed using random primers. Double-stranded cDNAs were end-repaired and 3’ adenylated. Adaptors were ligated at the 3’-end of the cDNAs. Libraries were PCR amplified using rolling circle amplification to make DNA nanoballs (DNB) and sequenced in multiplex as 100 bp paired-end reads. Adapters were trimmed with the SOAPnuke v2.2.1 (104) and reads were filtered on base quality to reach an average Phred+33 score of 37. Quality control was conducted using FastQC (105) leading to an average of 60.2 M quality-filtered reads per replicate. Reads were then mapped onto the AALBF3.v2 assembly using STAR v2.7.10a (106) with default parameters, except a minimum intron size of 10 bp. Mapping quality was assessed with samtools v1.6 (http://www.htslib.org/), with an average of 94 % of reads being properly mapped (**S9 Table**). Gene read counts were quantified by STAR with default parameters.

#### Differential transcription

Differential transcription analysis was performed using DESeq2 (Love et al., 2014) on all genes and pseudogenes showing ≥5 read counts in each of the four replicates for at least one mosquito line. Differentially transcribed genes were identified based on their congruent transcription fold change (FC) and adjusted Student’ t-test *p* values (False Discovery Rate for multiple testing correction, 107) across the five ‘resistant vs susceptible’ comparisons (see experimental design on **Fig. 1**). Fold change thresholds were set as follows: FC ≥ 3 in ‘Sel vs NS’; FC ≥ 3 in ‘Sel vs SRun’ and FC ≥ 1.5 in the three ‘field vs SRun’ comparisons while adjusted *p* Value threshold was set to 0.001 for all comparisons. In order to clarify the role of detoxification genes over-transcribed in association with resistance, their closest homologs were identified in *Ae. aegypti* and *An. gambiae* using a protein blast against *Ae. aegypti* (LVP_AGWG) and *An. gambiae* (PEST) proteomes from VeuPathDB (108,109). As P450 gene descriptions and gene names can vary across different genome versions, blast results were backed up with a phylogenetic analysis performed on CYP6s protein sequences from *Ae. albopictus*, *Ae. aegypti* and *An. gambiae*. Protein sequences were retrieved from NCBI and those having a validated role in pyrethroid resistance were identified based on the literature and sequence alignments. Phylogenetic trees were built using the only ‘NGPhylogeny.fr’ pipeline using approximate likelihood-ratio test (aLRT) as branch support, with a *CYP4* sequence as outgroup (110).

#### Alternative splicing

Alternative splicing events and indels were identified from RNA-seq data using KisSplice v2.6.2 (111). This *de novo* splicing events predictor was run on each replicate of each line with default parameters, except the parameter –b which was set to 15 to account for the higher level of polymorphism in *Ae. albopictus* compared with mammals. Splicing variants were identified with KissDE R package v1.19.0 (112). Only variants covered by at least 20 reads, affecting known genes and showing a congruent splicing variation across all ‘resistant vs susceptible’ comparisons were retained. Delta Percent Spliced In (ΔPSI) threshold were as follows: 20% for ‘Sel vs NS’; 20% for ‘Sel vs SRun’ and 15% for the three ‘field vs SRun’ comparisons. The adjusted *p* value threshold was set to 0.05 (Benjamini & Hochberg FDR for multiple testing correction). The splicing events passing these filters were then mapped to AALBF3.v2 assembly using KisSplice2RefGenome v1.2.3 (https://kissplice.prabi.fr/tools/kiss2refgenome/) and assigned into three types: alternative splicing, indels and intron retentions. The distinction between genomic indels and intron retention was based on an arbitrary threshold (indel if ≤ 50 bp and intron retention if > 50 bp).

### Whole genome pool sequencing

For each line, three batches of 33-35 three days-old virgin non-blood-fed females were sampled for whole genome pool-sequencing (100 individuals per line). Genomic DNA was extracted separately for each batch using the GENTRA Puregene kit (QIAGEN) following manufacturer’s instructions quantified using Qubit (Qiagen) and then pooled in equal DNA quantity. After quality check, one sequencing library was prepared for each line following the DNBseq plant and animal whole genome resequencing protocol (https://en.mgi-tech.com/products/). Briefly, DNA was fragmented, size selected (insert size ∼350 bp) and end-repaired. Adapters were ligated to the DNA fragments which were then size-selected, amplified by PCR using rolling circle amplification and sequenced in multiplex as 150 bp paired-end reads with the DNBseq Technology (MGI Tech Co). Sequencing reads were filtered based on base quality resulting in a final average Phred score of 37 and adapters were removed with the SOAPnuke software v2.2.1 (104) leading to an average of ∼1630 M reads per line (**S9 Table**). Reads were then mapped onto the *Ae. albopictus* AALBF3.v2 genome with BWA mem v0.7.17-r1188 (113). SAM files were converted into BAM format, sorted by coordinates and unmapped reads were removed using samtools v 1.6 (114). Mapped reads were trimmed and only reads showing a probability of correct mapping of 0.999 were kept (MapQ ≥30). PCR duplicates were removed using picard MarkDuplicates v2.27.5 (http://broadinstitute.github.io/picard), resulting in an average of 631 M mapped reads per sample and a mean genome coverage of 64.2X.

#### Copy number variations

Genomic duplications were searched for by looking for genes showing a higher mean read coverage in resistant than in susceptible mosquito lines. Though likely to generate false positives, this approach was preferred to pipelines integrating local coverage variations and the presence of flanking non-colinear reads (115) because the *Ae. albopictus* is genome is highly enriched with transposable elements (>41% of the genome, 89) likely to generate false negatives. Indeed, transposable elements are frequent at duplication breakpoints leading to the loss of non-colinear reads (e.g. split reads and both mate flip reads) due to multiple mapping events. Since intronic regions are highly variable and thus more subject to local coverage variations, mean normalized gene coverages were computed from exonic regions only. Fold changes of normalized exonic coverage between lines were computed for all genes. Since very low gene coverage can lead to artefactual high coverage variations, genes showing a normalized coverage lower than the 5% global quantile were not considered. Furthermore, a global observation of the data set indicated that the dispersion of coverage variations increased for smaller transcripts. To take this into account, a sliding windows approach considering transcript length was applied to identify differentially covered genes: for each pairwise line comparison, a gene was considered as affected by a positive copy number variation (CNV) when its normalized coverage variation was higher than the 95% quantile of the variation observed from the 1000 genes showing the most similar transcript lengths. Following this approach, genes were considered as showing a positive CNV associated with resistance when passing this threshold for all ‘resistant vs susceptible’ comparisons (‘Sel vs NS’, ‘Sel vs SRun’ and the three ‘field vs SRun’ comparisons).

#### Gene polymorphisms

Gene polymorphisms associated with resistance were searched for using reads mapped to the coding regions of protein-coding genes. For each line, polymorphisms were called using bcftools-v1.17 (116). We then retained biallelic variants showing a read coverage between 50 and 300 reads in all lines, a frequency ≥ 15% in at least one line and a frequency variation ≥ 15% between at least two lines. Genic effects were annotated according to the longest reading frame using snpEff v5.2c (117). As the reference genome Foshan strain is susceptible to deltamethrin (118) variants were considered associated with resistance if the frequency of the variant allele increased in ‘resistant vs susceptible’ comparisons. Frequency thresholds were as follows: frequency ≤ 15% in the susceptible SRun line, frequency variation ≥ 25% in ‘Sel vs SRun’ and in ‘Sel vs NS’ comparisons, and frequency variation ≥ 20% in the three ‘Field vs SRun’ comparisons.

We also specifically searched for non-synonymous polymorphisms affecting the gene encoding the voltage-gated sodium channel protein targeted by deltamethrin (VGSC-para gene carrying the *Kdr* mutations). However, this gene was incorrectly assembled in AALBF3 likely due to uncontrolled heterozygosity. Indeed, the previous AALFBF2 assembly (Palatini et al., 2020) contains two annotated gene copies, each located in a different scaffold (the ‘VGSC-para’ LOC109421922, and the ‘VGSC para-like’ LOC109432678). In AALBF3, both genic sequences were present but the ‘para’ sequence was not annotated as a gene. In the latest AALBF5 assembly, a better control of heterozygosity led to the two contigs being merged, resulting in a single annotated ‘VGSC-para’ gene (LOC109421922). We therefore assumed that the AALBF5 assembly is correct. In order to study the polymorphism of this gene, all reads mapped to AALBF3.v2 in the para (Chr3:378648962-378932310) and para-like regions (Chr3:377089613-377411247) were re-mapped to the AALBF5 gene LOC109421922 and SNPs were called as described above.

#### Selection signatures

Polymorphisms data were also used to look for selection signatures associated with deltamethrin resistance between the Sel and NS lines, which only diverged by 5 generations of deltamethrin selection. This analysis was based on all exonic bi-allelic substitutions covered by 50 to 300 reads in each line and being polymorphic between these two lines (*i*.e. showing a variant frequency from 5% to 95% in at least one line). From this dataset, two genetic differentiation indexes were computed at the gene level: the mean Absolute Frequency Difference (mean AFD), and the percentage of SNPs in the top decile of the overall genomic AFD distribution. These indexes were plotted against each other and across the AALBF3.v2 genome for the 10192 protein-coding genes containing ≥ 20 substitutions.

## Acknowledgments

This study was funded by the French Agency for Food, Environmental and Occupational Health & Safety under the grant number EST-19-055. TB was supported by a PhD fellowship funded by the Grenoble-Alpes University (UGA). The funders had no role in study design, data collection and interpretation, or decision to submit the work for publication. We thank Dr. Julien Cattel for providing the susceptible reference strain SRun. We also thank Vincent Lefort for help with phylogenetic analyses.

## Data availability

RNA-seq and WG Pool-seq sequencing data have been deposited at EBI short read archive (SRA) under accession numbers E-MTAB-15418 and E-MTAB-15456.

## Ethical aspects

Mice were kept in the animal facility of the Biology department of the University of Grenoble-Alpes, approved by the French Ministry of Animal Welfare (agreement no. B 38 421 10 001) and used in accordance with the laws of the European Union (directive 2010/63/EU). The use of animals for this study was approved by the ComEth Grenoble-C2EA-12 ethics committee mandated by the French Ministry of Higher Education and Research (MENESR). The study was conducted in accordance with the ARRIVE guidelines.

## Author contributions

**Conceptualization**: Tiphaine Bacot, Jean-Marc Bonneville, Jean-Philippe David

**Data curation:** Tiphaine Bacot, Pascal Oberbach, Vincent Lacroix, Jean-Marc Bonneville

**Formal analysis**: Tiphaine Bacot, Pascal Oberbach, Jean-Marc Bonneville, Jean-Philippe David

**Funding acquisition**: Jean-Philippe David

**Investigation**: Tiphaine Bacot, Pascal Oberbach, Thierry Gaude, Louis Nadalin, Frederci Laporte, Guillaume Dupuy, Frederic Boyer

**Methodology**: Frederic Boyer, Vincent Lacroix, Tiphaine Bacot, Jean-Marc Bonneville, Jean-Philippe David

**Project administration**: Jean-Philippe David

**Resources**: Nausicaa Habchi-Hanriot, Guillaume Dupuy, Cyrille Czeher

**Supervision**: Jean-Marc Bonneville and Jean-Philippe David

**Writing – original draft**: Tiphaine Bacot, Vincent Lacroix, Jean-Marc Bonneville, Jean-Philippe David

**Writing – review & editing**: Tiphaine Bacot, Michael C Fontaine, Jean-Marc Bonneville, Jean-Philippe David

## Supporting information

**S1 File. Polymorphisms in the VGSC gene**. The first xlsx sheet describes all polymorphisms detected in the VGSC coding regions. For each polymorphism, the following information is provided: Position relative to AALBF5 assembly; amino-acid position in respect to the longest isoform; genic effect; variant allele frequency in each line; Variant allele frequency variations between resistant and susceptible lines; number of reads supporting each allele for each line. The second xlsx sheet provides a graphical overview of the position of the variant identified relative to transmembrane domains and known *Kdr* mutations.

**S2 Table. Gene transcription data**. This table describes transcription data for all genes detected by RNA-seq. For each gene the following information is provided: AALBF3 accession number; chromosomal position relative to AALBF3.v2 assembly; coding strand; gene type; Candidate gene family; baseMean (mean normalized read counts across all conditions); log_2_ fold change and adjusted *P* value for each resistant/susceptible comparison.

**S3 Table. Alternative splicing.** For each splicing event identified, the following information is provided: AALBF3 accession number; gene description; candidate gene family; chromosome; event start; event end; event type and subtype; event size in bp; frameshift occurrence; mean coverage; percent spliced in (PSI for each line); PSI variation (ΔPSI for each comparison).

**S4 Table. Gene copy number variations**. For each gene detected, the following information is provided: AALBF3 accession number; chromosomal position relative to the AALBF3.v2 assembly; coding strand; gene type; candidate gene family; gene description; mRNA length; normalized exonic coverage (for each lines); mean normalized coverage across all lines; coverage filter; CNV status for both lab and field comparisons.

**S5 Table. Non-synonymous variants associated with resistance**. For each variant, the following information is provided: chromosomal position relative to AALBF3.v2 assembly; reference and variant alleles; polymorphism type; AALBF3 gene accession number; gene description; candidate gene family; longest transcript; variant exon rank; variant genic effect; amino-acid change; cDNA position relative to cDNA length; amino-acid position relative to protein length; total coverage (for each line); variant allele frequency (for each line).

**S6 Table. Gene differentiation indexes**. For each gene the following information is provided: Gene accession number; chromosomal position relative to AALBF3.v2 assembly; coding strand; gene description; candidate gene family; total number of polymorphic SNPs; number of synonymous SNPs; number of non-synonymous SNPs; number of resistance-associated non-synonymous SNPs; mean Allele Frequency Variation between Sel and NS Lines (mean AFD); the percent of SNPs showing an AFD within the top decile of the global distribution (% top decil AFD); differential transcription associated with resistance (Yes/No); gene being within the inversion identified on chr2 (Yes/No).

**S7 Table. The AALBF3.v2 genome assembly**. The genomic position of AALBF3 scaffolds on AALBF3.v2 chromosomes is provided. AALBF3 scaffolds were assembled in the same order as described in Boyle et al. 2021 with 50N spacers inserted between them.

**S8 Table. Resistance candidate genes**. For each gene the following information is provided: AALBF3accesion number; chromosomal position relative to AALBF3.v2 assembly; coding strand; gene type; candidate gene family; gene description.

**S9 File. Sequencing data overview**. The first xlsx sheet provides RNA-seq sequencing and mapping statistics. The second xlsx sheet provides whole genome Pool-seq sequencing and mapping statistics.

## References

1. Orr HA. The genetic theory of adaptation: a brief history. Nat Rev Genet. 2005;6(2):119–27.

2. Hendry AP, Farrugia TJ, Kinnison MT. Human influences on rates of phenotypic change in wild animal populations. Mol Ecol. 2008;17(1):20–9.

3. Hendry AP, Gotanda KM, Svensson EI. Human influences on evolution, and the ecological and societal consequences. Philos Trans R Soc Lond B Biol Sci. 2017;372(1712).

4. Palumbi SR. Humans as the World’s Greatest Evolutionary Force. Science. 2001;293(5536):1786–90.

5. Ffrench-Constant RH, Daborn PJ, Le Goff G. The genetics and genomics of insecticide resistance. Trends Genet. 2004;20(3):163–70.

6. Liu. Insecticide Resistance in Mosquitoes: Impact, Mechanisms, and Research Directions. Annu Rev Entomol. 2015;60(1):537–59.

7. Bass C, Jones CM. Editorial overview: Pests and resistance: Resistance to pesticides in arthropod crop pests and disease vectors: mechanisms, models and tools. Curr Opin Insect Sci. 2018;27:iv–vii.

8. Dusfour I, Vontas J, David JP, Weetman D, Fonseca DM, Corbel V, et al. Management of insecticide resistance in the major Aedes vectors of arboviruses: Advances and challenges. PLoS Negl Trop Dis. 2019;13(10):e0007615.

9. Hawkins NJ, Bass C, Dixon A, Neve P. The evolutionary origins of pesticide resistance. Biol Rev Camb Philos Soc. 2018;94(1):135–155.

10. Li XC, Schuler MA, Berenbaum MR. Molecular mechanisms of metabolic resistance to synthetic and natural xenobiotics. Annu Rev Entomol. 2007;52:231–53.

11. Brown JE, Evans BR, Zheng W, Obas V, Barrera-Martinez L, Egizi A, et al. Human impacts have shaped historical and recent evolution in Aedes aegypti, the dengue and yellow fever mosquito. Evolution. 2014;68(2):514–25.

12. Paupy C, Delatte H, Bagny L, Corbel V, Fontenille D. Aedes albopictus, an arbovirus vector: from the darkness to the light. Microbes Infect Inst Pasteur. 2009;11(14-15):1177–85.

13. Bonizzoni M, Gasperi G, Chen X, James AA. The invasive mosquito species Aedes albopictus: current knowledge and future perspectives. Trends Parasitol. 2013;29(9):460–8.

14. Carvalho VL, Long MT. Perspectives on New Vaccines against Arboviruses Using Insect-Specific Viruses as Platforms. Vaccines. 2021;9(3):263.

15. Garg H, Mehmetoglu-Gurbuz T, Joshi A. Virus Like Particles (VLP) as multivalent vaccine candidate against Chikungunya, Japanese Encephalitis, Yellow Fever and Zika Virus. Sci Rep. 2020;10(1):4017.

16. Silva JVJ, Lopes TRR, Oliveira-Filho EFD, Oliveira RAS, Durães-Carvalho R, Gil LHVG. Current status, challenges and perspectives in the development of vaccines against yellow fever, dengue, Zika and chikungunya viruses. Acta Trop. 2018;182:257–63.

17. Moyes CL, Vontas J, Martins AJ, Ng LC, Koou SY, Dusfour I, et al. Contemporary status of insecticide resistance in the major Aedes vectors of arboviruses infecting humans. PLoS Negl Trop Dis. 2017;11(7):e0005625.

18. Van Den Berg H, Da Silva Bezerra HS, Al-Eryani S, Chanda E, Nagpal BN, Knox TB, et al. Recent trends in global insecticide use for disease vector control and potential implications for resistance management. Sci Rep. 2021;11(1):23867.

19. Dusfour I, Thalmensy V, Gaborit P, Issaly J, Carinci R, Girod R. Multiple insecticide resistance in Aedes aegypti (Diptera: Culicidae) populations compromises the effectiveness of dengue vector control in French Guiana. Mem Inst Oswaldo Cruz. 2011;106(3):346–52.

20. Marcombe S, Carron A, Darriet F, Etienne M, Agnew P, Tolosa M, et al. Reduced efficacy of pyrethroid space sprays for dengue control in an area of Martinique with pyrethroid resistance. Am J Trop Med Hyg. 2009;80(5):745–51.

21. Marcombe S, Darriet F, Tolosa M, Agnew P, Duchon S, Etienne M, et al. Pyrethroid resistance reduces the efficacy of space sprays for dengue control on the island of Martinique (Caribbean). PLoS Negl Trop Dis. 2011;5(6):e1202.

22. Valle D, Bellinato DF, Viana-Medeiros PF, Lima JBP, Martins Junior ADJ. Resistance to temephos and deltamethrin in Aedes aegypti from Brazil between 1985 and 2017. Mem Inst Oswaldo Cruz. 2019;114:e180544.

23. Smith LB, Kasai S, Scott JG. Pyrethroid resistance in Aedes aegypti and Aedes albopictus: Important mosquito vectors of human diseases. Pestic Biochem Physiol. 2016;133:1–12.

24. Soderlund DM. Molecular mechanisms of pyrethroid insecticide neurotoxicity: recent advances. Arch Toxicol. 2012;86(2):165–81.

25. Auteri M, La Russa F, Blanda V, Torina A. Insecticide Resistance Associated with kdr Mutations in Aedes albopictusJJ: An Update on Worldwide Evidences. BioMed Res Int. 2018;2018:1–10.

26. Endersby-Harshman NM, Schmidt TL, Hoffmann AA. Diversity and distribution of sodium channel mutations in Aedes albopictus (Diptera: Culicidae). Healy K, éditeur. J Med Entomol. 2024;61(3):630–43.

27. Soderlund DM, Knipple DC. The molecular biology of knockdown resistance to pyrethroid insecticides. Insect Biochem Mol Biol. 2003;33(6):563–77.

28. Uemura N, Itokawa K, Komagata O, Kasai S. Recent advances in the study of knockdown resistance (kdr) mutations in Aedes mosquitoes with a focus on several remarkable mutations. Curr Opin Insect Sci. 2024;101178.

9. Chambers J. An introduction to the metabolism of pyrethroids. In: Gunther FA, Gunther JD, éditeurs. Residue Reviews [Internet]. New York, NY: Springer New York; 1980. p. 101–24. (Reviews of Environmental Contamination and Toxicology; vol. 73). http://link.springer.com/10.1007/978-1-4612-6068-4_7

30. Dermauw W, Van Leeuwen T. The ABC gene family in arthropods: Comparative genomics and role in insecticide transport and resistance. Insect Biochem Mol Biol. 2014;45:89–110.

31. Panini M, Manicardi GC, Moores GD, Mazzoni E. An overview of the main pathways of metabolic resistance in insects. Invertebr Surviv J. 2016;326–335 Pages.

32. Faucon F, Dusfour I, Gaude T, Navratil V, Boyer F, Chandre F, et al. Identifying genomic changes associated with insecticide resistance in the dengue mosquito Aedes aegypti by deep targeted sequencing. Genome Res. 2015;25(9):1347–59.

33. Ffrench-Constant RH. The molecular genetics of insecticide resistance. Genetics. 2013;194(4):807–15.

34. Lucas ER, Miles A, Harding NJ, Clarkson CS, Lawniczak MK, Kwiatkowski DP, et al. Whole genome sequencing reveals high complexity of copy number variation at insecticide resistance loci in malaria mosquitoes. BioRxiv. 2018;10.1101/39956.

35. Weetman D, Djogbenou LS, Lucas E. Copy number variation (CNV) and insecticide resistance in mosquitoes: evolving knowledge or an evolving problem? Curr Opin Insect Sci. 2018;27:82–8.

36. Chuaycharoensuk T, Juntarajumnong W, Boonyuan W, Bangs MJ, Akratanakul P, Thammapalo S, et al. Frequency of pyrethroid resistance in Aedes aegypti and Aedes albopictus (Diptera: Culicidae) in Thailand. J Vector Ecol. 2011;36(1):204–12.

37. Ishak HI, Zairi J, Ranson H, Wondji C. Contrasting patterns of insecticide resistance and knockdown resistance (kdr) in the dengue vectors Aedes aegypti and Aedes albopictus from Malaysia. Parasit Vectors. 2015;8:181.

38. Lee RML, Choong CTH, Goh BPL, Ng LC, Lam-Phua SG. Bioassay and biochemical studies of the status of pirimiphos-methyl and cypermethrin resistance in Aedes (Stegomyia) aegypti and Aedes (Stegomyia) albopictus (Diptera: Culicidae) in Singapore. Trop Biomed. 2014;31(4):670–9.

39. Thanispong K, Sathantriphop S, Malaithong N, Bangs MJ, Chareonviriyaphap T. Establishment of Diagnostic Doses of Five Pyrethroids for Monitoring Physiological Resistance in Aedes Albopictus in Thailand. J Am Mosq Control Assoc. 2015;31(4):346–52.

40. Gao JP, Chen HM, Shi H, Peng H, Ma YJ. Correlation between adult pyrethroid resistance and knockdown resistance (kdr) mutations in Aedes albopictus (Diptera: Culicidae) field populations in China. Infect Dis Poverty. 2018;7(1):86.

41. Kushwah RBS, Mallick PK, Ravikumar H, Dev V, Kapoor N, Adak TP, et al. Status of DDT and pyrethroid resistance in Indian Aedes albopictus and absence of knockdown resistance (kdr) mutation. J Vector Borne Dis. 2015;52(1):95–8.

42. Mohsin M, Naz S, Khan I, Jabeen A, Bilal H, Ahmed R, et al. Susceptibility status of Aedes aegypti and Aedes albopictus against insecticides at eastern Punjab, Pakistan. Int J Mosq Res. 2016;3:41–6.

43. Kasai S, Caputo B, Tsunoda T, Cuong TC, Maekawa Y, Lam-Phua SG, et al. First detection of a Vssc allele V1016G conferring a high level of insecticide resistance in Aedes albopictus collected from Europe (Italy) and Asia (Vietnam), 2016: a new emerging threat to controlling arboviral diseases. Euro Surveill Bull Eur Sur Mal Transm Eur Commun Dis Bull. 2019;24(5).

44. Zhou X, Yang C, Liu N, Li M, Tong Y, Zeng X, et al. Knockdown resistance (kdr) mutations within seventeen field populations of Aedes albopictus from Beijing China: first report of a novel V1016G mutation and evolutionary origins of kdr haplotypes. Parasit Vectors. 2019;12(1):180.

45. Ishak IH, Riveron JM, Ibrahim SS, Stott R, Longbottom J, Irving H, et al. The Cytochrome P450 gene CYP6P12 confers pyrethroid resistance in kdr-free Malaysian populations of the dengue vector Aedes albopictus. Sci Rep. 2016;6:24707.

46. Marcombe S, Doeurk B, Thammavong P, Veseli T, Heafield C, Mills MA, et al. Metabolic Resistance and Not Voltage-Gated Sodium Channel Gene Mutation Is Associated with Pyrethroid Resistance of Aedes albopictus (Skuse, 1894) from Cambodia. Insects. 2024;15(5):358.

47. Xu J, Su X, Bonizzoni M, Zhong D, Li Y, Zhou G, et al. Comparative transcriptome analysis and RNA interference reveal CYP6A8 and SNPs related to pyrethroid resistance in Aedes albopictus. Attardo GM, éditeur. PLoS Negl Trop Dis. 2018;12(11):e0006828.

48. Pichler V, Bellini R, Veronesi R, Arnoldi D, Rizzoli A, Lia RP, et al. First evidence of resistance to pyrethroid insecticides in Italian AEDES ALBOPICTUS populations 26 years after invasion. Pest Manag Sci. 2018;74(6):1319–27.

49. Pichler V, Malandruccolo C, Paola S, Bellini R, Severini F, Toma L, et al. Phenotypic and genotypic pyrethroid resistance of Aedes albopictus, with focus on the 2017 chikungunya outbreak in Italy. Pest Manag Sci. 2019;

50. Pichler V, Caputo B, Valadas V, Micocci M, Horvath C, Virgillito C, et al. Geographic distribution of the V1016G knockdown resistance mutation in Aedes albopictus: a warning bell for Europe. Parasit Vectors. 2022;15(1):280.

51. Djiappi-Tchamen B, Nana-Ndjangwo MS, Mavridis K, Talipouo A, Nchoutpouen E, Makoudjou I, et al. Analyses of Insecticide Resistance Genes in Aedes aegypti and Aedes albopictus Mosquito Populations from Cameroon. Genes. 2021;12(6):828.

52. Yamashita S, Uruma K, Yang C, Higa Y, Minakawa N, Cuamba N, et al. The origin and insecticide resistance of Aedes albopictus mosquitoes established in southern Mozambique. Parasit Vectors. 2024;17(1):292.

53. Yougang AP, Keumeni CR, Wilson-Bahun TA, Tedjou AN, Njiokou F, Wondji C, et al. Spatial distribution and insecticide resistance profile of Aedes aegypti and Aedes albopictus in Douala, the most important city of Cameroon. Plos One. 2022;17(12):e0278779.

54. Delatte H, Dehecq JS, Thiria J, Domerg C, Paupy C, Fontenille D. Geographic Distribution and Developmental Sites of Aedes albopictus (Diptera: Culicidae) During a Chikungunya Epidemic Event. Vector-Borne Zoonotic Dis. 2008;8(1):25–34.

55. Delatte H, Paupy C, Dehecq JS, Thiria J, Failloux AB, Fontenille D. Aedes albopictus, vecteur des virus du chikungunya et de la dengue à la RéunionJJ: biologie et contrôle. Parasite. 2008;15(1):3–13.

56. Renault P, Solet JL, Sissoko D, Balleydier E, Larrieu S, Filleul L, et al. A major epidemic of chikungunya virus infection on Reunion Island, France, 2005-2006. Am J Trop Med Hyg. 2007;77(4):727–31.

57. Vincent M, Paty MC, Gerardin P, Balleydier E, Etienne A, Daoudi J, et al. From dengue outbreaks to endemicity: Reunion Island, France, 2018 to 2021. Eurosurveillance. 2023;28(29).

58. Expertise from the French National Agency for Health, Food and Environmental Safety (ANSES) n°2020-SA-0029. Résistance des moustiques vecteurs aux insecticides. 2020. https://www.anses.fr/fr/system/files/BIOCIDES2020SA0029Ra.pdf

59. Tancredi A, Papandrea D, Marconcini M, Carballar-Lejarazu R, Casas-Martinez M, Lo E, et al. Tracing temporal and geographic distribution of resistance to pyrethroids in the arboviral vector Aedes albopictus. Aldridge RL, éditeur. PLoS Negl Trop Dis. 2020;14(6):e0008350.

60. Chandor-Proust A, Bibby J, Regent-Kloeckner M, Roux J, Guittard-Crilat E, Poupardin R, et al. The central role of mosquito cytochrome P450 CYP6Zs in insecticide detoxification revealed by functional expression and structural modelling. Biochem J. 2013;455:75–85.

61. Nikou D, Ranson H, Hemingway J. An adult-specific CYP6P450 gene is overexpressed in a pyrethroid-resistant strain of the malaria vector, Anopheles gambiae. Gene. 2003;318:91–102.

62. Vontas J, Katsavou E, Mavridis K. Cytochrome P450-based metabolic insecticide resistance in Anopheles and Aedes mosquito vectors: Muddying the waters. Pestic Biochem Physiol. 2020;170:104666.

63. Stevenson BJ, Bibby J, Pignatelli P, Muangnoicharoen S, O’Neill PM, Lian LY, et al. Cytochrome P450 6M2 from the malaria vector Anopheles gambiae metabolizes pyrethroids: Sequential metabolism of deltamethrin revealed. Insect Biochem Mol Biol. 2011;41(7):492–502.

64. Kouamo MFM, Ibrahim SS, Hearn J, Riveron JM, Kusimo M, Tchouakui M, et al. Genome-Wide Transcriptional Analysis and Functional Validation Linked a Cluster of Epsilon Glutathione S-Transferases with Insecticide Resistance in the Major Malaria Vector Anopheles funestus across Africa. Genes. 2021;12(4):561.

65. Messenger LA, Impoinvil LM, Derilus D, Yewhalaw D, Irish S, Lenhart A. A whole transcriptomic approach provides novel insights into the molecular basis of organophosphate and pyrethroid resistance in Anopheles arabiensis from Ethiopia. Insect Biochem Mol Biol. 2021;139:103655.

66. Simma EA, Dermauw W, Balabanidou V, Snoeck S, Bryon A, Clark RM, et al. Genome-wide gene expression profiling reveals that cuticle alterations and P450 detoxification are associated with deltamethrin and DDT resistance in Anopheles arabiensis populations from Ethiopia. Pest Manag Sci. 2019;75(7):1808–18.

67. Tao F, Si F, Hong R, He X, Li X, Qiao L, et al. Glutathione S –transferase (GST) genes and their function associated with pyrethroid resistance in the malaria vector Anopheles sinensis. Pest Manag Sci. 2022;78(10):4127–39.

68. Bariami V, Jones CM, Poupardin R, Vontas J, Ranson H. Gene Amplification, ABC Transporters and Cytochrome P450s: Unraveling the Molecular Basis of Pyrethroid Resistance in the Dengue Vector, Aedes aegypti. PLoS Negl Trop Dis. 2012;6(6).

69. Bonizzoni M, Afrane Y, Dunn WA, Atieli FK, Zhou G, Zhong D, et al. Comparative transcriptome analyses of deltamethrin-resistant and –susceptible Anopheles gambiae mosquitoes from Kenya by RNA-Seq. PloS One. 2012;7(9):e44607.

70. Epis S, Porretta D, Mastrantonio V, Urbanelli S, Sassera D, De Marco L, et al. Temporal dynamics of the ABC transporter response to insecticide treatment: insights from the malaria vector Anopheles stephensi. Sci Rep. 2014;4(1):7435.

71. He Q, Yan Z, Si F, Zhou Y, Fu W, Chen B. ATP-Binding Cassette (ABC) Transporter Genes Involved in Pyrethroid Resistance in the Malaria Vector Anopheles sinensis: Genome-Wide Identification, Characteristics, Phylogenetics, and Expression Profile. Int J Mol Sci. 2019;20(6):1409.

72. Mastrantonio V, Ferrari M, Epis S, Negri A, Scuccimarra G, Montagna M, et al. Gene expression modulation of ABC transporter genes in response to permethrin in adults of the mosquito malaria vector Anopheles stephensi. Acta Trop. 2017;171:37–43.

73. Xu J, Zheng J, Zhang R, Wang H, Du J, Li J, et al. Identification and functional analysis of ABC transporter genes related to deltamethrin resistance in Culex pipiens pallens. Pest Manag Sci. 2023;79(10):3642–55.

74. Scheuer T. Regulation of sodium channel activity by phosphorylation. Semin Cell Dev Biol. 2011;22(2):160–5.

75. Srinivasan Y, Elmer L, Davis J, Bennett V, Angelides K. Ankyrin and spectrin associate with voltage-dependent sodium channels in brain. Nature. 1988;333(6169):177–80.

76. Manni M, Guglielmino CR, Scolari F, Vega-Rua A, Failloux AB, Somboon P, et al. Genetic evidence for a worldwide chaotic dispersion pattern of the arbovirus vector, Aedes albopictus. PLoS Negl Trop Dis. 2017;11(1):e0005332.

77. Maynard AJ, Ambrose L, Cooper RD, Chow WK, Davis JB, Muzari MO, et al. Tiger on the prowl: Invasion history and spatio-temporal genetic structure of the Asian tiger mosquito Aedes albopictus (Skuse 1894) in the Indo-Pacific. Vasilakis N, éditeur. PLoS Negl Trop Dis. 2017;11(4):e0005546.

78. Mousson L, Dauga C, Garrigues T, Schaffner F, Vazeille M, Failloux AB. Phylogeography of Aedes (Stegomyia) aegypti (L.) and Aedes (Stegomyia) albopictus (Skuse) (Diptera: Culicidae) based on mitochondrial DNA variations. Genet Res. 2005;86(1):1–11.

79. Chen H, Li K, Wang X, Yang X, Lin Y, Cai F, et al. First identification of kdr allele F1534S in VGSC gene and its association with resistance to pyrethroid insecticides in Aedes albopictus populations from Haikou City, Hainan Island, China. Infect Poverty. 2016;5:31.

80. David JP, Ismail HM, Chandor-Proust A, Paine MJ. Role of cytochrome P450s in insecticide resistance: impact on the control of mosquito-borne diseases and use of insecticides on Earth. Philos Trans R Soc Lond B Biol Sci. 2013;368(1612):20120429.

81. Bengoa M, Eritja R, Delacour S, Miranda MA, Sureda A, Lucientes J. First Data on Resistance to Pyrethroids in Wild Populations of Aedes albopictus from Spain. J Am Mosq Control Assoc. 2017;33(3):246–9.

82. Chen HM. Role of metabolic detoxification enzyme activity and knockdown resistance gene mutations in resistance of aedes albopictus to pyrethroid insecticides. Acad J Second Mil Med Univ. 2019;512–9.

83. Khan HAA. Resistance to insecticides and synergism by enzyme inhibitors in Aedes albopictus from Punjab, Pakistan. Sci Rep. 2020;10(1):21034.

84. Li Y, Xu J, Zhong D, Zhang H, Yang W, Zhou G, et al. Evidence for multiple-insecticide resistance in urban Aedes albopictus populations in southern China. Parasit Vectors. 2018;11(1):4.

85. Feyereisen R. Insect P450 enzymes. Annu Rev Entomol. 1999;44:507–33.

86. Bacot T, Haberkorn C, Guilliet J, Cattel J, Kefi M, Nadalin L, et al. A genomic duplication spanning multiple P450s contributes to insecticide resistance in the dengue mosquito Aedes aegypti. Peer Community J. 2024;4:e110.

87. Cattel J, Haberkorn C, Laporte F, Gaude T, Cumer T, Renaud J, et al. A genomic amplification affecting a carboxylesterase gene cluster confers organophosphate resistance in the mosquito Aedes aegyptiJJ: From genomic characterization to high-throughput field detection. Evol Appl. 2021;14(4):1009–22.

88. Remnant EJ, Good RT, Schmidt JM, Lumb C, Robin C, Daborn PJ, et al. Gene duplication in the major insecticide target site, Rdl, in Drosophila melanogaster. Proc Natl Acad Sci. 2013;110(36):14705–10.

89. Melo ESD, Wallau GL. Mosquito genomes are frequently invaded by transposable elements through horizontal transfer. Malik HS, éditeur. Plos Genet. 2020;16(11):e1008946.

90. Nene V, Wortman JR, Lawson D, Haas B, Kodira C, Tu ZJ, et al. Genome sequence of Aedes aegypti, a major arbovirus vector. Science. 2007;316(5832):1718–23.

91. Palatini U, Masri RA, Cosme LV, Koren S, Thibaud-Nissen F, Biedler JK, et al. Improved reference genome of the arboviral vector Aedes albopictus. Genome Biol. 2020;21(1):215.

92. Amezian D, Nauen R, Le Goff G. Transcriptional regulation of xenobiotic detoxification genes in insects – An overview. Pestic Biochem Physiol. 2021;174:104822.

93. Ji M, Vandenhole M, De Beer B, De Rouck S, Villacis-Perez E, Feyereisen R, et al. A nuclear receptor HR96-related gene underlies large trans-driven differences in detoxification gene expression in a generalist herbivore. Nat Commun. 2023;14(1):4990.

94. Wen X, Feng K, Qin J, Wei P, Cao P, Zhang Y, et al. A detoxification pathway initiated by a nuclear receptor TcHR96h in Tetranychus cinnabarinus (Boisduval). Palli SR, éditeur. Plos Genet. 2023;19(9):e1010911.

95. Lien NTK, Ngoc NTH, Lan NN, Hien NT, Tung NV, Ngan NTT, et al. Transcriptome Sequencing and Analysis of Changes Associated with Insecticide Resistance in the Dengue Mosquito (Aedes aegypti) in Vietnam. Am J Trop Med Hyg. 2019;100(5):1240–8.

96. Shan G, Hammock BD. Development of Sensitive Esterase Assays Based on α-Cyano-Containing Esters. Anal Biochem. 2001;299(1):54–62.

97. Faria R, Johannesson K, Butlin RK, Westram AM. Evolving Inversions. Trends Ecol Evol. 2019;34(3):239–48.

98. Haberkorn C, David J, Henri H, Delpuech J, Lasseur R, Vavre F, et al. A major 6 Mb superlocus is involved in pyrethroid resistance in the common bed bug Cimex lectularius. Evol Appl. 2023;16(5):1012–28.

99. Wellenreuther M, Mérot C, Berdan E, Bernatchez L. Going beyond SNPs: The role of structural genomic variants in adaptive evolution and species diversification. Mol Ecol. 2019;28(6):1203–9.

100. Wellenreuther M, Bernatchez L. Eco-Evolutionary Genomics of Chromosomal Inversions. Trends Ecol Evol. 2018;33(6):427–40.

101. Deng J, Guo Y, Su X, Liu S, Yang W, Wu Y, et al. Impact of deltamethrin-resistance in Aedes albopictus on its fitness cost and vector competence. Churcher TS, éditeur. PLoS Negl Trop Dis. 2021;15(4):e0009391.

102. Standard operating procedure for testing insecticide susceptibility of adult mosquitoes in WHO tube tests. SOP version: WHO Tube test/01/14. World Health Organization; 2022. https://iris.who.int/bitstream/handle/10665/352316/9789240043831-eng.pdf?sequence=1

103. Boyle JH, Rastas PMA, Huang X, Garner AG, Vythilingam I, Armbruster PA. A Linkage-Based Genome Assembly for the Mosquito Aedes albopictus and Identification of Chromosomal Regions Affecting Diapause. Insects. 2021;12(2):167.

104. Chen Y, Chen Y, Shi C, Huang Z, Zhang Y, Li S, et al. SOAPnuke: a MapReduce acceleration-supported software for integrated quality control and preprocessing of high-throughput sequencing data. GigaScience. 2018;7(1):1–6.

105. Andrews S. FastQC: a quality control tool for high throughput sequence data. 2010. Available online at: http://www.bioinformatics.babraham.ac.uk/projects/fastqc/

106. Dobin A, Davis CA, Schlesinger F, Drenkow J, Zaleski C, Jha S, et al. STAR: ultrafast universal RNA-seq aligner. Bioinformatics. 2013;29(1):15–21.

107. Benjamini Y, Hochberg Y. Controlling the False Discovery Rate: a Practical and Powerful Approach to Multiple Testing. J R Stat Soc B. 1995;57:289–300.

108. Giraldo-Calderón GI, Harb OS, Kelly SA, Rund SS, Roos DS, McDowell MA. VectorBase.org updates: bioinformatic resources for invertebrate vectors of human pathogens and related organisms. Curr Opin Insect Sci. 2022;50:100860.

109. Harb OS, McDowell MA, Roos DS. VEuPathDB Resources: A Platform for Free Online Data Exploration, Integration, and Analysis. In: Setubal JC, Stadler PF, Stoye J, editors. Comparative Genomics. New York, NY: Springer US; 2024 p. 573–86. (Methods in Molecular Biology; vol. 2802).

110. Lemoine F, Correia D, Lefort V, Doppelt-Azeroual O, Mareuil F, Cohen-Boulakia S, et al. NGPhylogeny.fr: new generation phylogenetic services for non-specialists. Nucleic Acids Res. 2019;47(W1):W260-5.

111. Sacomoto GA, Kielbassa J, Chikhi R, Uricaru R, Antoniou P, Sagot MF, et al. KIS SPLICE: de-novo calling alternative splicing events from RNA-seq data. BMC Bioinformatics. 2012;13(S6):S5.

112. Benoit-Pilven C, Marchet C, Chautard E, Lima L, Lambert MP, Sacomoto G, et al. Complementarity of assembly-first and mapping-first approaches for alternative splicing annotation and differential analysis from RNAseq data. Sci Rep. 2018;8(1):4307.

113. Li H, Durbin R. Fast and accurate short read alignment with Burrows-Wheeler transform. Bioinformatics. 2009;25(14):1754–60.

114. Li H, Handsaker B, Wysoker A, Fennell T, Ruan J, Homer N, et al. The Sequence Alignment/Map format and SAMtools. Bioinforma Oxf Engl. 2009;25(16):2078–9.

115. Schrider DR, Begun DJ, Hahn MW. Detecting highly differentiated copy-number variants from pooled population sequencing. Pac Symp Biocomput Pac Symp Biocomput. 2013;344–55.

116. Danecek P, Bonfield JK, Liddle J, Marshall J, Ohan V, Pollard MO, et al. Twelve years of SAMtools and BCFtools. Gigascience. 2021;10(2):giab008.

117. Cingolani P, Platts A, Wang LL, Coon M, Nguyen T, Wang L, et al. A program for annotating and predicting the effects of single nucleotide polymorphisms, SnpEff: SNPs in the genome of Drosophila melanogaster strain w JJ; iso-2; iso-3. Fly. 2012;6(2):80–92.

118. Lin L qun, Chen Y hui, Tian Y fan, Chen Y sen, Zheng Z yang, Wu J xin, et al. Study on the cross-resistance of Aedes albopictus (Skuse) (Diptera: Culicidae) to deltamethrin and pyriproxyfen. Parasit Vectors. 2024;17(1):403.

